# Relationships between aboveground plant traits and carbon cycling in tundra plant communities

**DOI:** 10.1101/865899

**Authors:** Konsta Happonen, Anna-Maria Virkkala, Julia Kemppinen, Pekka Niittynen, Miska Luoto

## Abstract

1. The functional composition and diversity of plant communities are globally applicable predictors of ecosystem functioning. Yet, it is unclear how traits influence carbon cycling. This is an important question in the tundra where vegetation shifts are occurring across the entire biome, and where soil organic carbon stocks are large and vulnerable to environmental change.
2. To study how traits affect carbon cycling in the tundra, we built a model that explained carbon cycling (above-ground and soil organic carbon stocks, and photosynthetic and respiratory fluxes) with abiotic conditions (air temperature and soil moisture), plant community functional composition (average plant height, leaf dry matter content (LDMC) and specific leaf area (SLA)), and functional diversity (weighted standard deviations of the traits). Data was collected from an observational study setting from northern Finland.
3. The explanatory power of the models was relatively high, but a large part of variation in soil organic carbon stocks remained unexplained. Plant height was the strongest predictor of all carbon cycling variables except soil carbon stocks. Communities of larger plants were associated with larger CO_2_ fluxes and above-ground carbon stocks. Communities with fast leaf economics (i.e. high SLA and low LDMC) had higher photosynthesis, ecosystem respiration, and soil organic carbon stocks.
4. Within-community variability in plant height, SLA, and LDMC affected ecosystem functions differently. SLA and LDMC diversity increased CO_2_ fluxes and soil organic carbon stocks, while height diversity increased the above-ground carbon stock. The contributions of functional diversity metrics to ecosystem functioning were about as important as those of average SLA and LDMC traits.
5. Synthesis: Plant height, SLA, and LDMC have clear effects on tundra carbon cycling. The importance of functional diversity highlights a potentially important mechanism controlling the vast tundra carbon pools that should be better recognized. More research on root traits and decomposer communities is needed to understand the below-ground mechanisms regulating carbon cycling in the tundra.

## Introduction

The Arctic is warming two to four times faster than the world on average (Holland and Landrum 2015; Post et al. 2019), leading to changes in plant growth (Myers-Smith et al. 2015), height (Bjorkman et al. 2018), and species distributions (Steinbauer et al. 2018). It is unclear how such changes in vegetation will influence the carbon stocks and balance of these ecosystems. On one hand, the expansion of larger plants might increase net carbon uptake (Cahoon et al. 2012; Sørensen et al. 2019). On the other hand, vegetation-driven changes in soil conditions and microorganisms might accelerate decomposition and, in turn, increase carbon losses to the atmosphere (Parker, Subke, and Wookey 2015; Vowles and Björk 2018). Arctic soils contain more than half of the global soil organic carbon stock (Hugelius et al. 2014), thus changes in tundra vegetation and carbon cycling have the potential to influence atmospheric CO_2_ concentrations at the global scale.

Plant functional traits provide a quantitative and globally applicable means to study the interactions among vegetation and carbon cycling in the tundra (Thomas et al. 2020). Above-ground traits of plant species and the functional composition of plant communities have been found to vary primarily along two axes (Díaz et al. 2016; Bruelheide et al. 2018). The first axis describes the trade-offs between resource acquisitive (fast) and conservative (slow) strategies, and has often been called the leaf economics spectrum (Wright et al. 2004) that is measured with traits such as specific leaf area (SLA), leaf nitrogen content, and leaf dry matter content (LDMC) (Bruelheide et al. 2018). The other axis is related to plant size, characterizing the trade-offs between investments to light competition, and photosynthesis and reproduction, and can be measured with, for example, plant vegetative height or stem specific density (Díaz et al. 2016). The plant size axis, and in particular plant height, has been observed to increase with climate change over the past decades across the tundra, whereas changes in leaf economic spectrum have been more variable (Bjorkman et al. 2018).

These trait axes explain differences between species’ vital rates that define population processes (Adler et al. 2014), and ecosystem functioning such as carbon cycling from local to global scales (Diaz et al. 2004; Michaletz et al. 2014). In the tundra, photosynthesis has been shown to correlate positively with fast leaf economics, indicated by, for example, high average SLA (Sørensen et al. 2019). This is because fast communities have more photosynthetic machinery relative to leaf area compared to slower communities (Shipley et al. 2006; Wright et al. 2004). Communities with fast traits often have higher ecosystem respiration as well because maintaining photosynthetic capacity is costly (Cavaleri, Oberbauer, and Ryan 2008). Plant size is often described using biomass or its proxies, such as total plant cover or leaf area index (LAI) (Oberbauer et al. 2007; Street et al. 2007; Shaver et al. 2007; Marushchak et al. 2013). Studies have shown that larger size leads to higher photosynthesis, net carbon uptake, and sometimes also soil organic carbon stocks (Lafleur and Humphreys 2018; Gagnon, Domine, and Boudreau 2019).

The relationships of leaf economic and plant size-related traits with soil organic carbon stocks can be more complex. This is because these stocks might be more indirectly linked to traits, via, for example, the quantity and quality of litter inputs (DeMarco, Mack, and Bret-Harte 2014), by affecting microbial communities or soil microclimate (Parker, Subke, and Wookey 2015; Kemppinen et al. 2021), or by correlating with traits that more directly affect soil processes, such as root chemical composition (G. T. Freschet et al. 2010; Bergmann et al. 2020). Earlier research indicates that faster communities such as meadows often accumulate large amounts of carbon into the soil whereas shrubs, and in particular deciduous shrub communities have smaller soil organic carbon stocks and high soil respiration (Sørensen et al. 2018; Vowles and Björk 2018; Parker, Subke, and Wookey 2015). However, the effects of above-ground traits of plant species on carbon cycling have not yet been comprehensively studied.

Climate change-driven changes in tundra vegetation are predicted to influence the diversity of communities (Niittynen, Heikkinen, and Luoto 2020) which also influences carbon cycling (Tilman, Isbell, and Cowles 2014; Duffy, Godwin, and Cardinale 2017). In the tundra, some evidence points out that more diverse shrub communities located in transition zones might be less productive than communities dominated by one shrub species due to competition (Fletcher et al. 2012). However, in general, the effects of tundra biodiversity on carbon cycling remain poorly understood. Using functional traits allows partitioning the diversity effect to contributions from complementarity along different niche axes and helps to understand how carbon cycling is influenced by changes in functional diversity, which represents one important component of biodiversity (Cadotte, Carscadden, and Mirotchnick 2011).

In this study, we examine the effect of average plant height, SLA, and LDMC and the within-community variability in these traits on growing season ecosystem CO_2_ fluxes and above-ground and soil organic carbon stocks in a tundra ecosystem (Table 1). We use a systematic study setting of up to 129 intensively measured plots and regression models while controlling for the effects of key microclimate variables of soil moisture and air temperature explaining carbon cycling (Poyatos et al. 2014; Nobrega and Grogan 2008; Siewert 2018; López-Blanco et al. 2017).

**Table 1.**
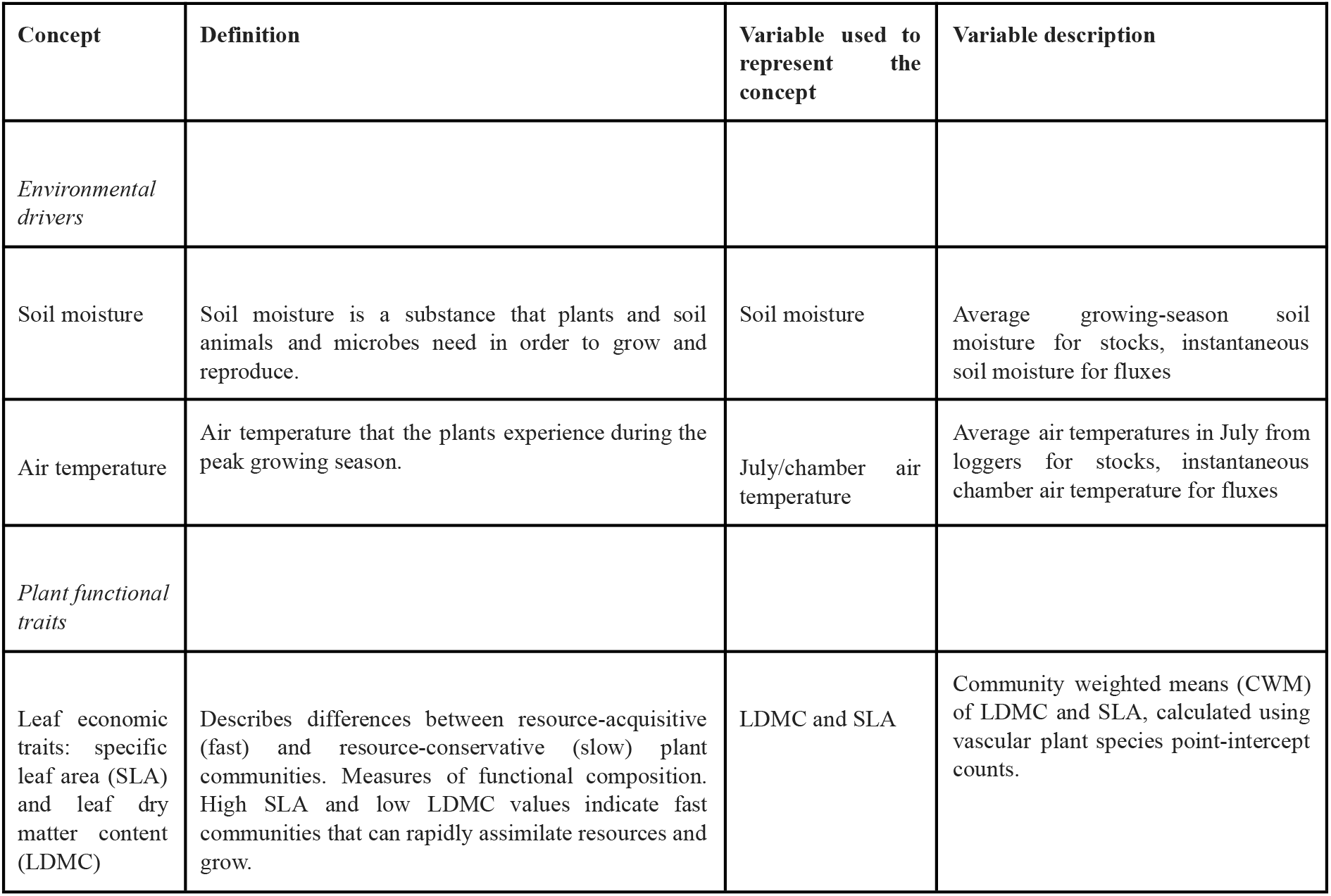
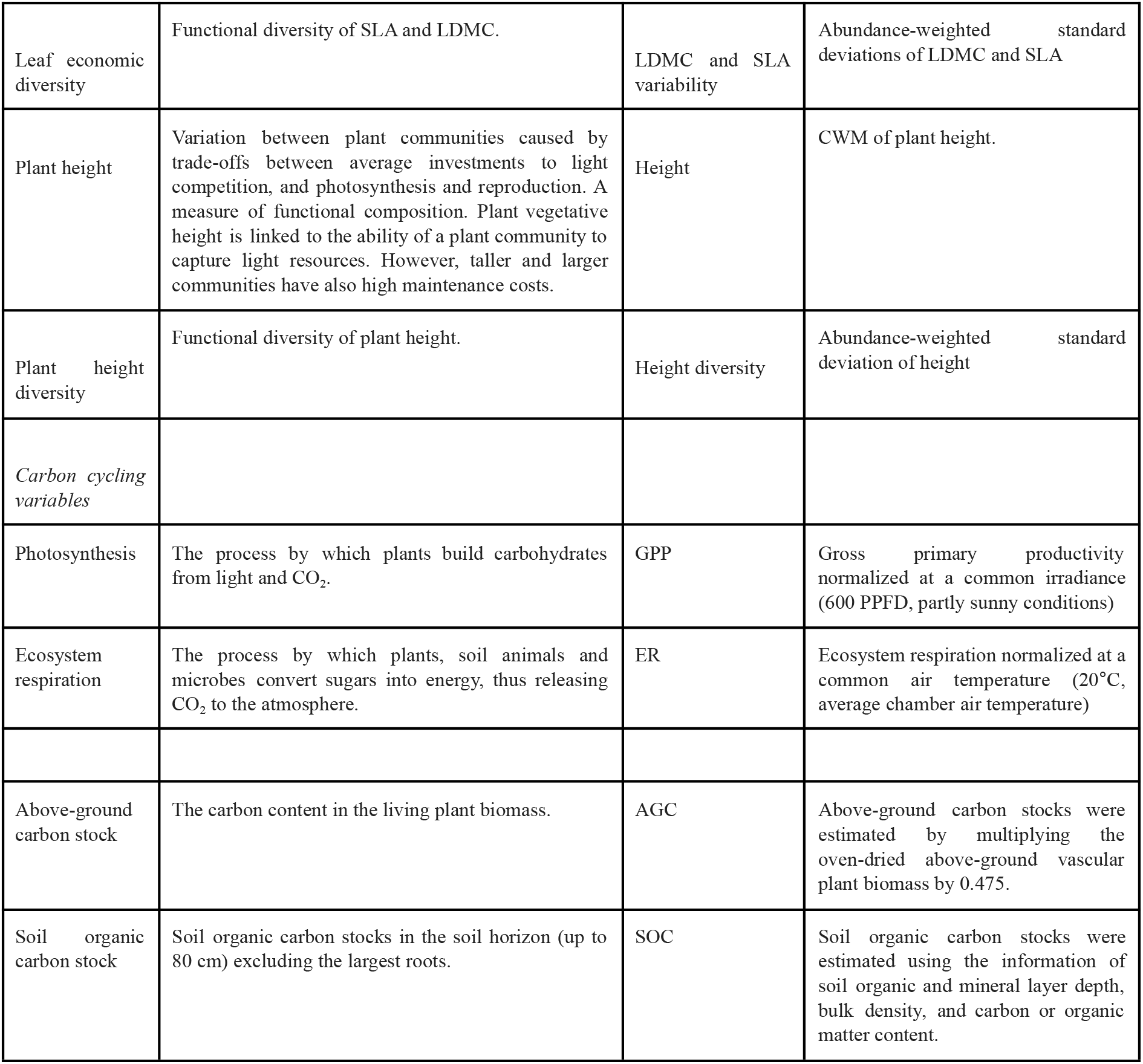
Glossary

| Concept | Definition | Variable used to represent the concept | Variable description |
| --- | --- | --- | --- |
| *Environmental drivers* |  |  |  |
| Soil moisture | Soil moisture is a substance that plants and soil animals and microbes need in order to grow and reproduce. | Soil moisture | Average growing-season soil moisture for stocks, instantaneous soil moisture for fluxes |
| Air temperature | Air temperature that the plants experience during the peak growing season. | July/chamber air temperature | Average air temperatures in July from loggers for stocks, instantaneous chamber air temperature for fluxes |
| *Plant functional traits* |  |  |  |
| Leaf economic traits: specific leaf area (SLA) and leaf dry matter content (LDMC) | Describes differences between resource-acquisitive (fast) and resource-conservative (slow) plant communities. Measures of functional composition. High SLA and low LDMC values indicate fast communities that can rapidly assimilate resources and grow. | LDMC and SLA | Community weighted means (CWM) of LDMC and SLA, calculated using vascular plant species point-intercept counts. |
| Leaf economic diversity | Functional diversity of SLA and LDMC. | LDMC and SLA variability | Abundance-weighted standard deviations of LDMC and SLA |
| Plant height | Variation between plant communities caused by trade-offs between average investments to light competition, and photosynthesis and reproduction. A measure of functional composition. Plant vegetative height is linked to the ability of a plant community to capture light resources. However, taller and larger communities have also high maintenance costs. | Height | CWM of plant height. |
| Plant height diversity | Functional diversity of plant height. | Height diversity | Abundance-weighted standard deviation of height |
| *Carbon cycling variables* |  |  |  |
| Photosynthesis | The process by which plants build carbohydrates from light and CO<sub>2</sub>. | GPP | Gross primary productivity normalized at a common irradiance (600 PPFD, partly sunny conditions) |
| Ecosystem respiration | The process by which plants, soil animals and microbes convert sugars into energy, thus releasing CO<sub>2</sub> to the atmosphere. | ER | Ecosystem respiration normalized at a common air temperature (20°C, average chamber air temperature) |
| Above-ground carbon stock | The carbon content in the living plant biomass. | AGC | Above-ground carbon stocks were estimated by multiplying the oven-dried above-ground vascular plant biomass by 0.475. |
| Soil organic carbon stock | Soil organic carbon stocks in the soil horizon (up to 80 cm) excluding the largest roots. | SOC | Soil organic carbon stocks were estimated using the information of soil organic and mineral layer depth, bulk density, and carbon or organic matter content. |

## Materials and Methods

### Study site

The field observations were collected in 2016–2018 in a subarctic tundra environment in Kilpisjärvi, northwestern Finland (Fig. 1). The study area is located on an elevational gradient between two mountains, Saana (1029 m.a.s.l) and Korkea-Jehkats (960 m.a.s.l), and the valley in between (ca. 600 m.a.s.l.). The study area is above the mountain birch (*Betula pubescens* var. pumila (L.) Govaerts) forest line, and is predominantly dwarf-shrub heath, with graminoid- and herb-rich meadows concentrated around meltwater streams. *Empetrum nigrum* L., *Betula nana* L., *Vaccinium myrtillus* L., *Vaccinium vitis-idaea* L. and *Phyllodoce caeruleae* (L.) Bab. are highly abundant in the area with graminoids and herbs such as *Deschampsia flexuosa* (L.) Trin. and *Viola biflora* L. dominant along the streams. The main herbivores in the area are reindeer (*Rangifer tarandus tarandus* L.), and voles and lemmings (Cricetidae). The soils in the area are mostly poorly developed leptosoils with shallow organic layers and occasional podzolization; however, the meadows have soils with thicker organic layers. The mean annual air temperature and annual precipitation at a nearby meteorological station (Kilpisjärvi kyläkeskus, 480 m.a.s.l, 1.5 km from the study area, 1981–2010) are −1.9°C and 487 mm, respectively (Pirinen et al. 2012). The annual air temperature was −0.4°C in 2016, −1.5°C in 2017, and −0.9°C in 2018, and precipitation 552 mm in 2016, 551 mm in 2017, and 405 mm in 2018 (Finnish Meteorological Institute 2021).

**Figure 1.**
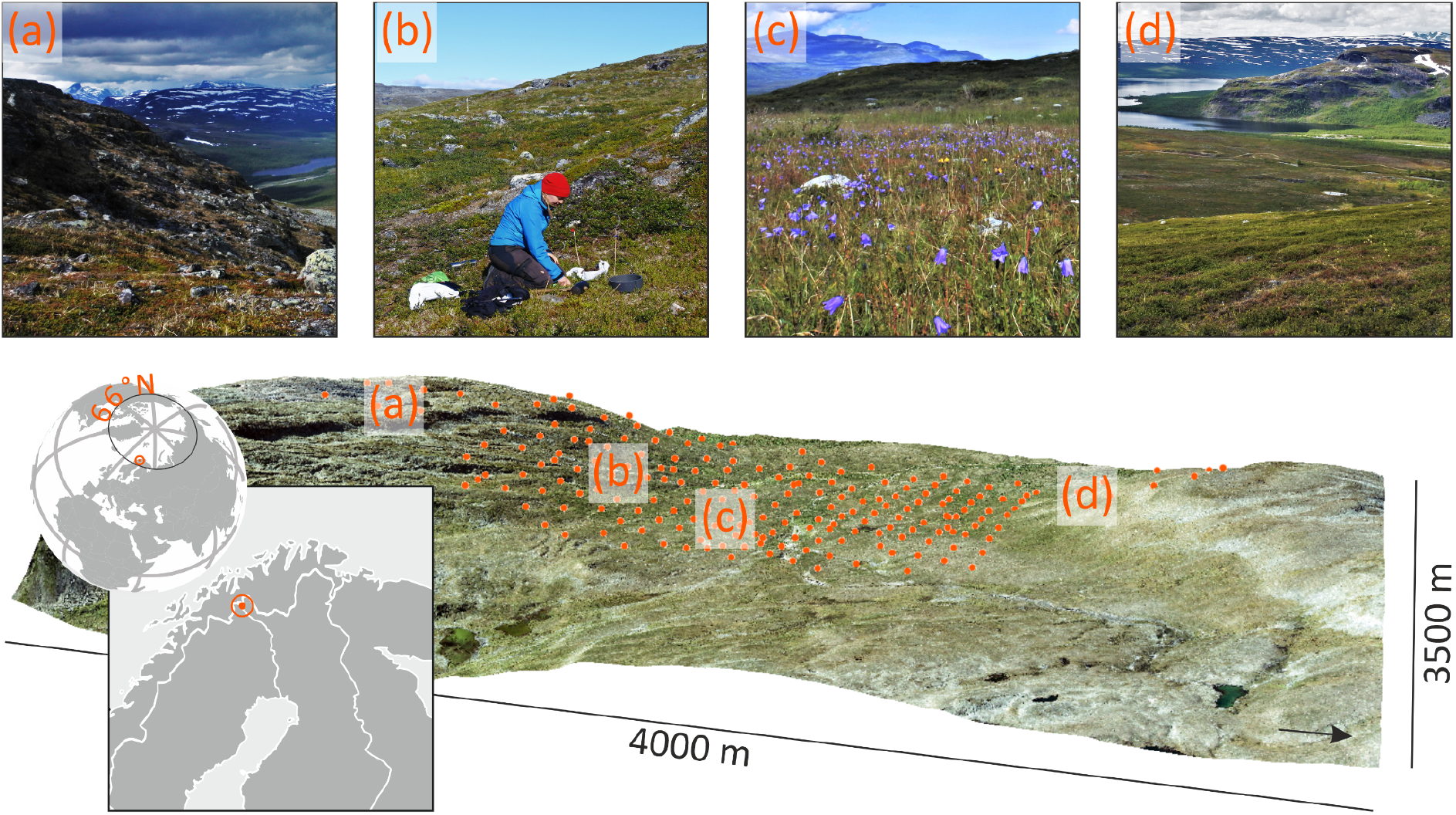
The study site. The vegetation is heterogenous. On the mountaintops and rocky slopes the vegetation consists of short species that can have either fast or slow leaf economic traits (a). Intermediately productive sites vary in vegetation height, and can be dominated by dwarf-shrubs with varying leaf economic strategies, such as deciduous *Vaccinium myrtillus* and *Betula nana* (b) or the evergreen *Empetrum nigrum* (d), with graminoids as subordinate species. The communities with tall shrubs can have a high variability in plant height. Lush and productive meadows can be found in the valley bottom. They consist of tall herbs and graminoids with high SLA and low LDMC, and often have high variability in both of these traits (c). The orthophoto (0.5 m resolution) is provided by the National Land Survey of Finland under a Creative Commons Attribution 4.0 licence (https://www.maanmittauslaitos.fi/en/maps-and-spatial-data/expert-users/product-descriptions/orthophotos, accessed on 2019-10-28.). Photographs by Julia Kemppinen.

The study design consisted of 129 locations in a 1.5 × 3.0 km area (Fig. 1). The distance between two adjacent locations was a minimum of 23 m (average 101 m). Individual study locations were thus in separate vegetation patches. Not all variables were measured in all locations, and the number of replicates for each analysis varied between 117 and 129 (Table 2). At the sampling locations where all variables were collected, organic and mineral layer depths and longer-term (i.e. a three-year record of) soil moisture and air temperature were measured at the central plot of the measurement scheme (Fig. 2). Plant functional traits, above-ground carbon stocks, and CO_2_ fluxes measured together with instantaneous chamber air temperature and soil moisture were measured in another plot 1–3 meters away from the central plot to avoid perturbing these permanent plots; however, we made sure that the environmental conditions of this plot were similar to the central plot to guarantee that these plots are comparable with each other (see Fig. 2). Finally, soil samples were collected from the immediate vicinity of the trait-carbon plot. To ease reading, the different measurement instruments are listed separately in Table S1.

**Table 2.** Medians, median absolute deviations (MAD), sample sizes and values for carbon cycling variables and their predictors. Bayesian R^2^ values for GAMs explaining carbon cycling variables are listed as well. GPP=gross primary productivity normalized at a common irradiance, ER = ecosystem respiration normalized at a common air temperature, AGC = above-ground carbon stock, SOC = soil organic carbon stock. Diversities are abundance-weighted standard deviations.

| Variable | Median | MAD | Unit | N | R<sup>2</sup> |
| --- | --- | --- | --- | --- | --- |
| **Responses** |  |  |  |  |  |
| GPP | 4.53 | 1.39 | $\mu\text{mol CO}_2 \text{ m}^{-2} \text{ s}^{-1}$ | 129 | 0.52 |
| ER | 2.87 | 0.71 | $\mu\text{mol CO}_2 \text{ m}^{-2} \text{ s}^{-1}$ | 129 | 0.47 |
| AGC | 0.12 | 0.07 | $\text{kg C m}^{-2}$ | 117 | 0.52 |
| SOC | 4.73 | 1.93 | $\text{kg C m}^{-2}$ | 117 | 0.28 |
| **Environment** |  |  |  |  |  |
| July air temperature | 13.89 | 0.29 | $^{\circ}\text{C}$ | 117 | |
| Chamber air temperature | 12.27 | 2.05 | $^{\circ}\text{C}$ | 129 | |
| Growing-season moisture | 18.82 | 2.90 | VWC (%) | 117 |  |
| Chamber soil moisture | 20.97 | 5.00 | VWC (%) | 129 |  |
| **Traits** |  |  |  |  |  |
| Height | 6.63 | 1.97 | cm | 129 |  |
| Height diversity | 2.72 | 0.74 | cm | 129 |  |
| LDMC | 0.44 | 0.06 | unitless | 129 |  |
| LDMC diversity | 0.08 | 0.03 | unitless | 129 |  |
| SLA | 11.86 | 3.10 | $\text{mm}^2 \text{ mg}^{-1}$ | 129 | |
| SLA diversity | 5.72 | 2.05 | $\text{mm}^2 \text{ mg}^{-1}$ | 129 | |

**Figure 2.**
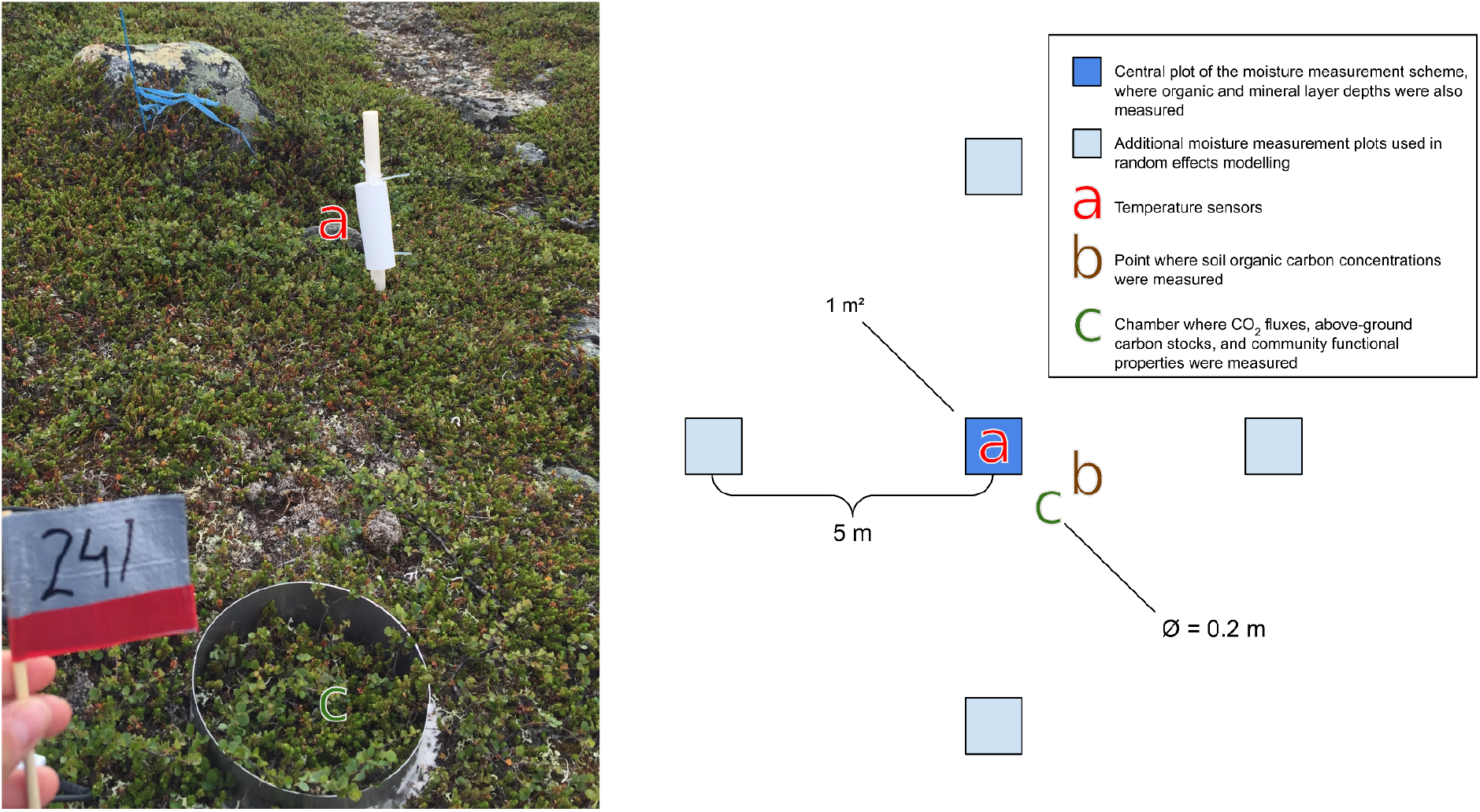
The measurement scheme. The image on the left displays a typical distance between the long-term temperature and moisture measurements (a) that were used to model carbon stocks, and the CO_2_ flux chamber (c), whose instantaneous temperature and soil moisture measurements were used to model CO_2_ fluxes. Soil organic carbon stocks were measured by combining information on carbon concentrations from point b with information on soil layer depths from point a.

### Environmental predictors of tundra carbon cycle variables

#### Microclimate variables: air temperature and soil moisture

Soil moisture was measured during the growing seasons of 2016–2018 3–6 times per year between June 6 and August 23 as volumetric water content (Happonen et al. 2019; Kemppinen et al. 2018). In each soil moisture plot, three measurements were taken, and the average was used in further analyses. Temperature loggers were installed in 2016 to monitor air temperature 10 cm above-ground at 2–4 h intervals. These air temperature measurements were aggregated to monthly averages. We used July air temperature in further analyses, because it is the warmest month in the area.

To quantify longer-term mean summer air temperature and soil moisture conditions (i.e., not representing just one year), we averaged July air temperature and growing season soil moisture levels over 2016–2018 for each location using random effects models with the R package lme4 (Bates et al. 2015). Air temperature were modelled using location and year as random effects (y ~ 1|location + year). In each location, soil moisture was measured from five 1 m^2^ plots: one central plot and one plot 5 m away in each cardinal direction to reflect fine-scale heterogeneity (Fig. 2). Soil moisture was modelled using plot nested in location, and year as random effects (y ~ 1|location/plot + year). Soil moisture was log-transformed before modelling. Finally, the environmental variables were predicted for the central plot of the soil moisture measurement scheme, averaging out yearly variation. In this landscape, soil moisture correlates positively and strongly with soil pH (see Happonen et al. 2019). Therefore, these soil moisture measurements partly also reflect the acidity of the soil, which might also be an indicator of soil nutrient concentrations.

Longer-term air temperature data were available for 86 plots with carbon stock measurements. To increase this number to 117, we interpolated average July air temperature using kriging with a spherical variogram model (R package gstat, Pebesma and Heuvelink 2016). Visual inspection revealed no effects of interpolation on the covariance and partial covariance structures of our data.

We further measured the air temperature inside the chamber and soil moisture directly next to the collar during the CO_2_ flux measurements to reflect instantaneous microclimate conditions during the flux measurement. We recorded soil moisture as the average of three measurements using the same devices as for the longer-term measurements; the air temperature measurements are described in section CO_2_ flux data.

#### Vegetation data: trait and community data

We quantified vascular plant community species composition from the plots where CO_2_ flux measurements were conducted using the point-intercept method during the peak growing season in 2017 (Happonen et al. 2019). A circular frame (20 cm diameter) with 20 evenly spaced pinholes was placed over the soil collar (see section CO_2_ flux data) after CO_2_ flux measurements. The total number of times each species touched pins (3 mm in thickness) lowered into the vegetation through the frame was counted and used as a measure of species abundance. While cryptogams might have important effects on soil processes, their biomass is considerably lower than that of vascular plants in subarctic heaths (M.C. Press et al. 1998). Furthermore, there is still no consensus on which quantitative traits of bryophytes and lichens best capture their effects on ecosystem functioning. For these reasons, the present study focuses on the effects of vascular plants on carbon cycling.

We measured plot-specific plant height, LDMC, and SLA for all species (Happonen et al. 2019). These three traits are among the most widely measured above-ground plant traits (Bruelheide et al. 2018; Díaz et al. 2016). Trait measurements were done separately for all locations, and always on plant individuals within the soil collars in order to account for intraspecific trait variation. Height was quantified as the height of the highest leaf on two random ramets within the collar. LDMC and SLA were measured from two leaf samples taken from two different ramets. The leaf samples were taken at the time of community surveys, put in re-sealable plastic bags with moist paper towels, and transported to the lab to be stored at 4°C for up to three days before scanning for leaf area and weighing for fresh mass. The leaves were then dried at 70°C for 48 h and weighed for dry mass. A precision scale with a resolution of 0.001 g was used for weighing. Location-specific trait values for each of the 70 observed vascular plant species were quantified as the average of the two individual trait values.

As a measure of functional composition, we used community weighted means (CWM) of these three traits following the mass-ratio hypothesis (Grime 1998), which states that ecosystem processes are determined by the trait values of dominant species in the community. CWMs of traits has been shown to correlate both with environmental conditions and ecosystem functioning (Garnier et al. 2004). We chose abundance-weighted standard deviation as the measure of within-community functional diversity. Functional composition and diversity were calculated separately for all traits.

### Carbon cycle variables

#### CO_2_ flux data

We measured CO_2_ exchange using a static, non-steady state non-flow-through system (Livingston and Hutchinson 1995) composed of a transparent acrylic chamber (20 cm diameter, 25 cm height). Measurements were done during the growing season as it is the most active season for plants with the largest net CO_2_ uptake (Belshe, Schuur, and Bolker 2013). Measurements were done in 2017 during which the weather conditions in the near-by meteorological station (10.8°C and 104 mm in July) were rather close to the average climatic conditions (11.2°C and 72 mm in July over 1981-2010). The chamber included a small ventilator, a carbon dioxide probe, an air humidity and temperature probe, and a measurement indicator. In the chamber, CO_2_ concentration and air temperature were recorded at 5-s intervals for 90 s. PAR was logged manually at 10-s intervals during the same period using a quantum sensor with a hand-held meter.

Steel soil collars (21 cm in diameter and 6–7 cm in height) were inserted in the soil at least 24 h before the measurements to avoid potential CO_2_ flush from soil due to the disturbance caused by the installation of the collars. The soils in the study area are relatively rocky and the plants have long horizontal roots, thus our collars could be embedded only *ca.* 2 cm into the soil. To guarantee an air-tight seal, we sealed the edges of the collar using inert quartz sand. The chamber was placed on top of the collar and ventilated after each measurement. We progressively decreased the light intensity of NEE measurements from ambient conditions to *ca.* 80%, 50% and 30% photosynthetic photon flux density (PPFD) by shading the chamber with layers of white mosquito net (n = 7–10). ER was measured in dark conditions (0 PPFD), which were obtained by covering the chamber with a space blanket (n = 3). Each plot was measured at midday during the peak season in 2017 between the 26^th^ of June and 27^th^ of July.

To convert concentration measurements into flux estimates, we deleted the first and last 5 s of each measurement series to remove potentially disturbed observations. The final measurement period was thus 80 seconds for each plot. Fluxes were calculated using linear regression and reported as μmol CO_2_ m^−2^ s^−1^. The median R^2^ of the flux estimates was 0.92. 10% of the fluxes had an R^2^ value below 0.25. Fluxes with a low R^2^ value were generally small (−0.5–0.5 μmol m^−2^ s^−1^) due to a low vegetation cover. These small fluxes were not removed from further analyses to guarantee that the plots cover larger abiotic and biotic environmental variability, and thus flux variability.

#### Carbon stock data

Stocks reflect the balance of carbon uptake and losses over a longer period of time. We measured the depth of the soil organic and mineral layers from three points on each central plot up to a depth of 80 cm using a soil corer. In the analyses, we used the mean value of the three point measurements to represent the organic and mineral layer depths in each sampling location. We collected samples of roughly 1 dl from the soil organic and mineral layers with metal soil core cylinders. The organic samples were collected from the top soil, and mineral samples directly below the organic layer. Organic samples were collected from all 117 plots and mineral samples from a subset of 51 plots. The soil samples were taken on the 1^st^ to 31^st^ of August 2016 and 2017, and were pre-processed and analysed at the Laboratory of Geosciences and Geography and the Laboratory of Forest Sciences, University of Helsinki.

Soil samples were first freeze-dried (ISO 11464:1994E). We analysed the bulk density from all organic samples, as well as the total carbon content (C%) or the soil organic matter content (SOM%). We estimated the bulk density (kg m^−3^) by dividing the dry weight by the sample volume. We used C% for the majority of the organic layer stock calculations, but for plots that were missing C% data we used SOM% (n = 24). We further analysed bulk density from 51 mineral samples and C% from 41 samples, respectively. We used the median bulk density and C% of these samples (820 kg m^−3^ for bulk density, range 470–1400 kg m^−3^; 3% for C%, range 0.4–6.5%) to estimate the carbon stock of the mineral layer. We used the median values, as we did not have data on the mineral soil conditions from all plots. We justified this approach since most of the soil carbon in this landscape is stored in the organic layer as the mineral soils are dry and have a coarse texture (see Kemppinen et al. 2018) that does not absorb organic matter particles. Before C% analysis, mineral samples were sieved through a 2 mm plastic sieve. Organic samples were homogenized by hammering the material into smaller pieces. We analysed C% using elemental analysers, and SOM% using the loss-on-ignition method according to SFS 3008.

SOC for organic and mineral layer was estimated using the following equations:

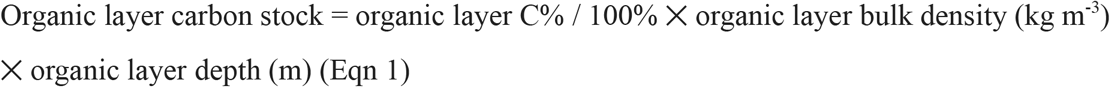

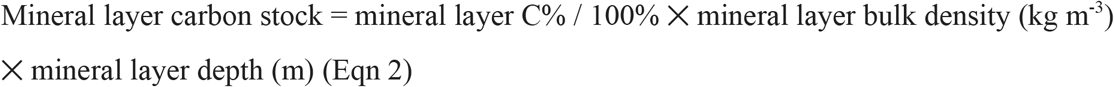

Finally, organic and mineral layer stocks were summed together to calculate the total SOC stock up to 80 cm. For plots that were missing organic layer C%, SOM% was used instead of C% in Eq. 1 to calculate the soil organic matter stocks, which we converted into organic layer carbon stocks based on a relationship between C% and SOM% (carbon fraction in the soil organic matter). We calculated the carbon fraction in the soil organic matter as 0.54 (R^2^ = 0.97), similar to Parker, Subke and Wookie (2015). For more details, see Kemppinen et al. (2021). Locations with no soil were excluded from the analysis.

Above-ground vascular plant biomass was collected between the 1^st^ and 10^th^ of August, 2017 from the collars. Biomass samples were oven-dried at 70°C for 48 h and weighed after drying. Above-ground carbon stocks (AGC) were estimated by multiplying the total biomass by 0.475 (Schlesinger, 1991).

### Statistical analyses

#### Light response model and flux normalization

All models were run in R version 3.5.3 with the package *brms* (Bürkner, 2018), which is an interface to the Bayesian modelling platform Stan (Carpenter *et al.*, 2017).

To account for the effects of variation in light levels and air temperature on CO_2_ fluxes, we fitted plot-specific light-response curves using a non-linear hierarchical bayesian model. We used the Michaelis-Menten equation to model the *n* instantaneous observations of NEE as a function of *k* plot-specific ER, maximum photosynthetic rate GPP_max_, and the half-saturation constant K (Eqns 1 & 5). ER, GPP_max_ and K were allowed to have plot-specific averages (Eqns 2–4 & 6–8), and ER also had an exponential air temperature (T) response (Eqn 2).

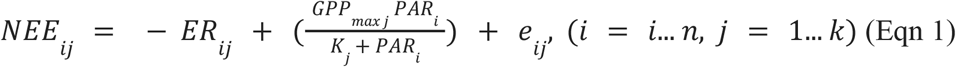

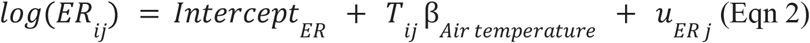

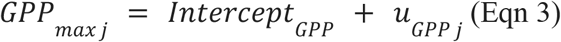

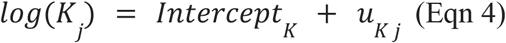

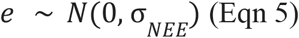

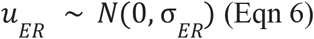

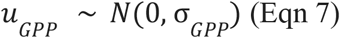

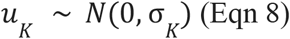

The Michaelis-Menten parameters GPP_max_ and half-saturation constant K sometimes identify weakly, so that the data would be consistent with infinitely increasing photosynthesis. This is especially true when CO_2_ fluxes are small, which is frequently the case in tundra ecosystems. To counter this, we set weakly informative priors on the plot-specific intercept terms based on visual inspection of the scale of variation in our data and typical parameter values reported in Williams *et al.*, (2006) (Eqns 9–11). A weakly informative prior was also set for the air temperature effect on respiration β_Air temperature_ (Eqn 12). Priors for the variance parameters were left as the weakly informative *brms* defaults (Eqn 13).

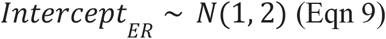

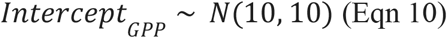

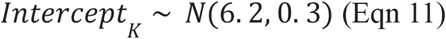

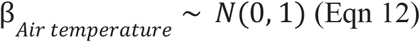

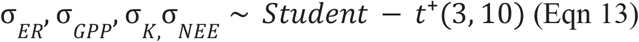

The model was fit with 4 Markov Chain Monte Carlo (MCMC) chains, which were run for 2000 iterations each. The first 1000 iterations were discarded as warmup, leaving a total of 4000 samples of each parameter.

We used this model to predict the rate of NEE at dark (0 PPFD) and average light (600 PPFD) conditions, and an air temperature of 20°C at each plot. 20°C was chosen as it corresponds to typical air temperature inside the chamber during flux measurements, and 600 PPFD because it is widely used in tundra literature (Street et al. 2007; Shaver et al. 2007; Dagg and Lafleur 2011; Marushchak et al. 2013)). Normalizing removes the effects of variation in instantaneous light and temperature conditions from the resulting variables, making them more comparable across the landscape. We refer to the flux normalized to dark conditions as ecosystem respiration (ER). We then subtracted ER from the NEE normalized to average light conditions to arrive at an estimate of normalized GPP.

#### Modeling tundra carbon cycling

We modelled GPP, ER, AGC, SOC, and organic layer SOC as functions of functional composition and functional diversity while adjusting for soil moisture and air temperature using bayesian generalized additive models (GAMs). All response variables were log-transformed before analyses to satisfy the assumption of homoscedasticity. Functional diversities were square root transformed to facilitate estimation of interactions by bringing very high values closer to the averages.

The effects of height and height diversity were included as thin-plate splines with basis dimension set to 15. A higher basis dimension allows estimation of wigglier responses, much in the same way that higher polynomials do (Wood 2017). Since LDMC and SLA are not independent, but both reflect variation along the leaf economic spectrum, the effects of leaf economic trait composition and diversity were modelled with tensor product smooths of their weighted averages and weighted standard deviations, respectively. A tensor product smooth allows estimation of wiggly and interpretable interaction surfaces that are much less rigid than interactions achieved using linear regression models (Pedersen et al. 2019). The thin plate basis dimension of each trait was set to five, resulting in a theoretical maximum degrees of freedom of 25 for the tensor product smooth. A similar tensor product smoother was also used to adjust for any remaining non-linear effects of air temperature, soil moisture and their interaction on CO_2_ fluxes and carbon stocks. Fluxes are controlled by instantaneous environmental conditions, whereas stocks are the balance of inputs and outputs over a longer period of time. For this reason, instantaneous air temperature and soil moisture measurements were used to adjust models of CO_2_ fluxes, while longer-term average air temperature and soil moisture were used for stocks.

Bayesian models require setting priors for the parameters. The intercept was given a student-t prior with an expectation of the empirical mean of the response, three degrees of freedom, and a standard deviation of 2.5. Each β parameter was given an uninformative flat prior over the reals. These are the default priors of the brms package. To reduce overfitting and excess wiggliness of the splines, we set the prior for the standard deviations of β parameters to an exponential distribution with a rate parameter of 1. All models were fitted with four MCMC chains of 2000 iterations. The first 1000 samples of each chain were discarded as warmup, leaving 4000 MCMC samples of each parameter of each model. The diagnostics for all bayesian models were satisfactory: no MCMC chain showed erratic behaviour under visual inspection, rank-normalized potential scale reduction factors for all parameters stayed below 1.01, and visual inspection of the posterior predictive distributions showed no large systematic differences between distributions of predicted and observed data. The median Bayesian R^2^ (Gelman et al. 2019) for the light-response model was 0.95. Bayesian R^2^ values for models of stocks and fluxes are presented in Table 2 and show that the model explaining soil organic carbon stocks had the lowest explanatory power.

CWMs and weighted standard deviations of SLA and LDMC are very correlated because their independent variation is restricted by trade-offs that underpin the leaf economics spectrum. This makes it somewhat unmeaningful to study responses in relation to just one leaf trait. Thus, to visualize the uncertainty in carbon cycling responses to both LDMC and SLA, and their variabilities, we post-processed our model predictions. First, using an errors-in-variables model (standard major axis regression, Legendre and Legendre 2012) written in Stan, we calculated an allometric relationship between LDMC and SLA, and LDMC variability and SLA variability. We then sampled values evenly from the lowest observed SLA value to the highest, and predicted the value of LDMC at this value of SLA. We then calculated the values of the tensor product smoothers at these trait values to visualize the uncertainty in carbon cycling responses to covarying SLA and LDMC. We repeated the procedure for the square root transformed leaf trait diversities.

Data and scripts used in this study are deposited into Zenodo (Happonen et al. 2021).

## Results

### Variability in environmental conditions and carbon cycle variables

Average values and within-landscape variabilities of carbon cycling, air temperature, soil moisture, traits and their diversities are presented in Table 2.

### Effects of air temperature and soil moisture on carbon cycling

Microclimate conditions and carbon cycle variables were in general positively linked to each other. In the light-response model, a one degree increase in air temperature resulted in a 2.9% (SE: 0.3%) increase in ER. In the GAM, air temperature and soil moisture increased GPP by 40% from the driest and coldest to the warmest and wettest parts of the gradient. AGC was predicted to be higher than average in warm and moist conditions, but lower than average in cool and moist environments. SOC stocks corresponded strongly with soil moisture and air temperature; average SOC stocks doubled along the gradient from cool and dry to warm and moist plots (Fig. 3). Organic layer SOC stocks correlated heavily with total SOC stocks (ρ=0.73, Fig. S1). Since organic layer SOC stocks are a subset of total SOC stocks and the results concerning these two stocks were qualitatively similar, results on organic SOC stocks are only presented in supplementary figures S2–S3.

**Figure 3.**
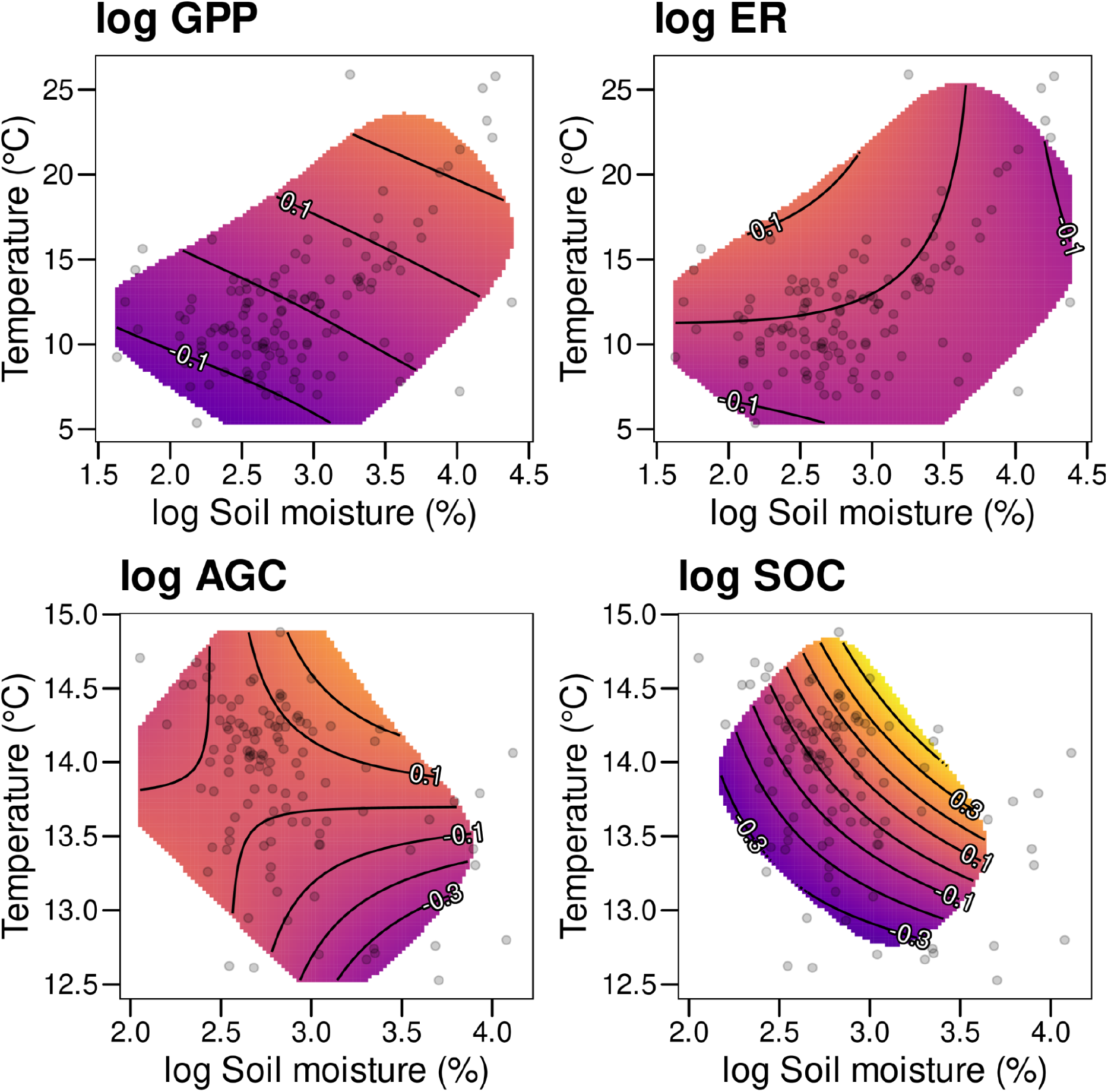
The marginal responses of CO_2_ fluxes and carbon stocks to the abiotic environment, modelled using a tensor product smooth of air temperature and soil moisture. For CO_2_ fluxes, the variables are chamber air temperature and soil moisture immediately next to the chamber during flux measurement. For carbon stock variables, the variables are 3-year average July air temperature and growing-season soil moisture. Point clouds show the observed data, and contour lines represent predicted relative responses on the log-scale (i.e. response variable on the z-axis). Responses are shown only where the standard error of the smoother is less than ¼ of the observed standard deviation. As respiration was already standardized to 20°C, it does not display an air temperature response here. A difference of 0.7 in the response represents about a 100% increase. GPP = Standardized gross primary productivity, ER = Standardized ecosystem respiration, AGC = Above-ground carbon stock, SOC = Soil organic carbon stock.

### Trait effects on CO_2_ fluxes

GPP and ER had very similar responses to plant community functional composition and diversity (Fig. 4). Fluxes increased with plant height, fast leaf traits (high SLA and low LDMC), and diversity in leaf traits. Variation in plant height had the largest absolute effect, as indicated by the larger range of marginal response values, with average GPP and ER increasing from shortest to tallest vegetation by 600% and 400%, respectively. Communities with the highest SLA and lowest LDMC had about 80% higher GPP and 50% higher ER compared to communities on the other end of this spectrum. Plots with the highest within-community variability in SLA and LDMC had about 80% higher GPP and ER compared to plots with the lowest variability.

**Figure 4.**
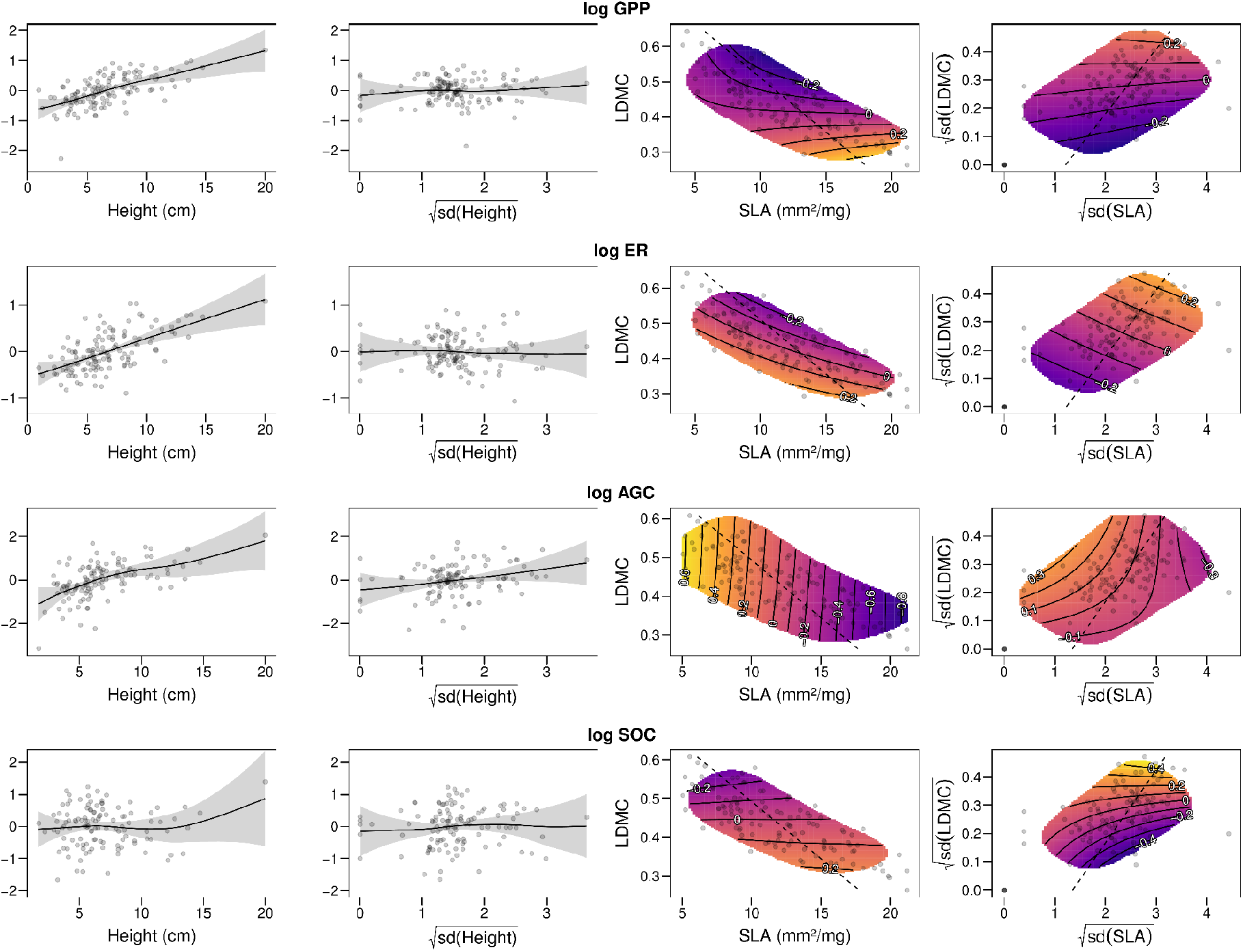
Marginal responses of CO_2_ fluxes (GPP = gross primary productivity normalized at a common irradiance, ER = ecosystem respiration normalized at a common air temperature), above-ground carbon stocks (AGC) and soil organic carbon stocks (SOC) to plant height, specific leaf area (SLA), leaf dry matter content (LDMC), and their within-community variability (standard deviations of the traits). In the two leftmost columns, the line represents the fitted response (i.e. response variable on the y-axis), the shading represents two standard errors and the points are partial residuals. In the two rightmost columns, point clouds show the observed data and contour lines represent predicted relative responses on the log-scale (i.e. response variable on the z-axis), and the dashed line shows the location of a model II regression line depicting the direction of covariation between SLA and LDMC, and their variabilities. The 2D response surfaces are limited to where the standard error of the smoother is less than 1/4 of the observed standard deviation. In all panels, a difference of 0.7 in the response represents about a 100% increase. In addition to the displayed variables, the models adjusted for the effects of instantaneous (fluxes) or long-term average (stocks) soil moisture, air temperature and their interaction (Fig. 3). Uncertainty in the responses of fluxes and stocks to LDMC, SLA, and their variabilities are depicted along the model II regression line in supplementary figures S4–8.

### Trait effects on carbon stocks

AGC increased with plant height and LDMC and decreased with SLA (Fig. 4). On the observed gradient of plant height, average AGC increased 23-fold. The gradient from low SLA and high LDMC to high SLA and low LDMC caused a ~70% reduction in AGC. There was also a 95% probability that AGC increases along with increasing height diversity, with the estimated difference in AGC between plots with the highest and lowest height diversity being about 4-fold on average.

Faster leaf economic traits (higher SLA and lower LDMC) and greater within-community variability in these leaf traits increased SOC. Along the gradient from low SLA - high LDMC communities to high SLA - low LDMC communities, SOC about doubled. Along the gradient from low variability in SLA and LDMC to high diversity in these leaf economic traits, SOC increased about 2.5-fold. We observed no clear links between SOC and plant height.

The effects of functional composition and diversity on carbon cycling are summarized in Fig. 5.

**Figure 5.**
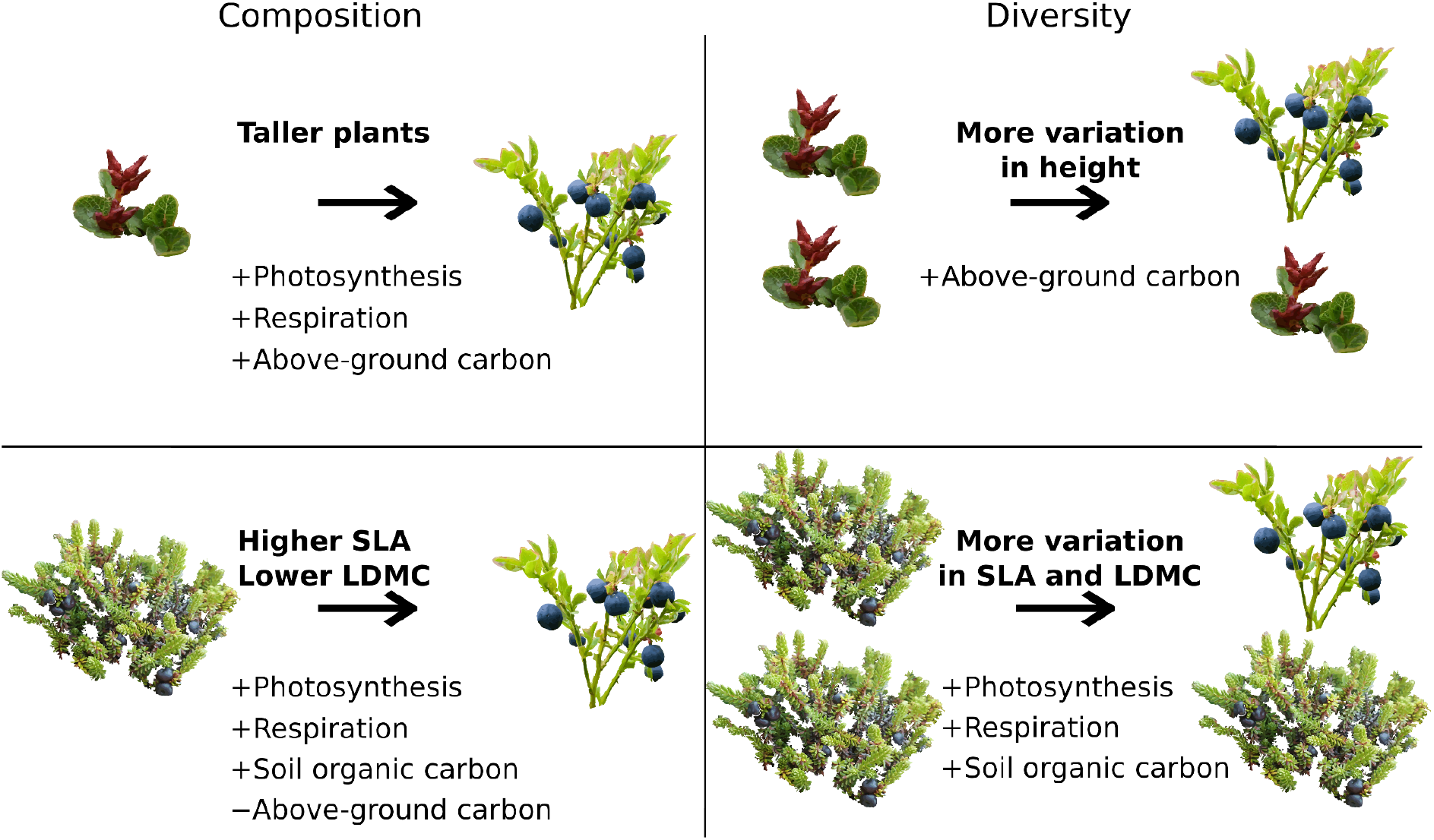
Summary of the relationships of peak-season CO_2_ fluxes and carbon stocks with the measures of functional composition and diversity used in this study, i.e. averages and within-community variabilities of plant height, specific leaf area (SLA) and leaf dry matter content (LDMC).

## Discussion

Arctic vegetation is reshuffled in various ways at an unprecedented speed (G. J. Jia, Epstein, and Walker 2003; Phoenix and Bjerke 2016; Post et al. 2009). Thus, it is imperative that we form a solid understanding of the causal forces shaping Arctic ecosystems and their functioning. Here, we show that a functional trait-based approach proved informative, at least according to the models in our observational study, since we were able to partition the effects of vegetation composition and diversity on carbon cycling to the effects of plant height, SLA, and LDMC, and their variability. Our results suggest that plant size and leaf economic traits have separate effects on tundra carbon cycling. Since these community trait attributes also respond differently to environmental conditions (Happonen et al. 2019; Bjorkman et al. 2018), they are well suited to inform us about the impacts of global change and vegetation shifts on carbon cycling and resulting climate feedbacks.

Our research further highlights the potential of using traits instead of plant functional groups or vegetation classifications which have often been used in earlier carbon cycle studies to describe the functional properties of vegetation (Virkkala et al. 2018). Using traits is advantageous as classifications might ignore large variation in vegetation properties that are relevant for ecosystem functioning (Thomas et al. 2019; Cadotte, Carscadden, and Mirotchnick 2011). Furthermore, the use of classification methods makes it impractical to compare the results of independent studies if their classifications differ whereas plant functional trait measurements are standardized and can be compared not only across the tundra but also across the globe.

Although we controlled for the key abiotic variables explaining functional composition, functional diversity, and carbon cycle variables, it is still possible that some confounding factors might be in part responsible for the relationships that we observed. Our study was further limited to a relatively warm and moderately rainy tundra region which does not represent the conditions of the entire biome but instead can be found in some parts of Fennoscandia, western Russia, Greenland, and Svalbard, in particular. The study area does not have permafrost, which covers most of the Arctic tundra (Brown et al. 2002), and has relatively small soil organic carbon stocks compared to the other parts of the tundra (Hugelius et al. 2014). Our results cannot thus be linked to how permafrost thaw might impact carbon cycling. And finally, our study did not cover the full spectrum of plant height present in the tundra discussed in the later sections of the Discussion. However, due to the local variability in environmental conditions, our study design covered almost all of the main vegetation types observed across the Arctic from barren lands to prostrate, evergreen and deciduous shrubs, and meadows, and large variability within these vegetation types (Fig. 1).

### Modeling the variability in the tundra carbon cycle

We observed high variability in CO_2_ fluxes and carbon stocks which was of similar magnitude as has been reported in other studies covering a range of environments across the entire Arctic during the growing season. For example, photosynthesis varied between 0.3 and 13 μmol CO_2_ m^−2^ s^−1^ in our study whereas those reported in Cahoon et al. (2012) in Greenland, Alaska, and Sweden varied from 1.5 to 10.2 (standardized similarly at an irradiance of 600 μmol CO_2_ m^−2^ s^−1^ of photosynthetic photon flux density). Observed soil organic carbon stocks (mostly 1–10 kg C m^−2^) were lower than those estimated for the upper 1 meter of the soil in this region by Hugelius et al., 2014 (5–15 kg m^−2^) but of similar magnitude as those reported for other Fennoscandian mountain regions (Sørensen et al. 2018; Ylänne et al. 2018). Nevertheless, observed soil organic carbon stocks were on average ca. 40 times larger than the above-ground carbon stocks, highlighting the fact that the soil carbon budget is highly important for the carbon balance of tundra ecosystems as a whole.

The models used in this study with R^2^ values of 0.28-0.52 had a similar or even higher explanatory power than what has been shown in earlier studies (e.g., models explaining photosynthesis had an R^2^ of 0.1-0.52 and models of ecosystem respiration 0.18-0.32 in Sørensen et al. (2019), models of photosynthesis had an R^2^ of 0.32 and of ecosystem respiration 0.45 in Mauritz et al. (2017)). Our light-response model showed that air temperature is important for respiration. In addition, soil moisture and air temperature had a positive relationship with photosynthesis and soil organic carbon stocks. This finding was similar to studies by López-Blanco et al. (2017), Nobrega and Grogan (2008), and Poytaos et al. (2014), that showed the widely-known importance of air temperature in controlling enzymatic processes, and soil moisture in acting as an important resource for organisms. We have also shown the importance of these variables on the distribution of plant functional traits in communities in our previous study: species with fast traits are favored in warmer and more moist parts of the landscape, while taller plants are found in warmer and drier conditions (Happonen et al. 2019). Above-ground carbon stocks were smaller in moist and cold environments, which might reflect a general scarcity of vascular plant vegetation in these environments.

### Plant height is the most important trait for most of the carbon cycle variables

Plant height had a strong and positive relationship with all growing season fluxes and above-ground carbon stocks, but not with soil organic carbon stocks. The relationship between plant height and photosynthesis was stronger than that of plant height and ecosystem respiration. Our study agrees with the results of Lafleur and Humphreys (2018), who discovered that taller plants (and in particular, shrubs) had larger growing season net CO_2_ uptake than lower plants in the low Arctic. They argued that this was likely due to colder soils limiting heterotrophic respiration and taller plants having a larger total leaf area to capture sunlight, which possibly also explains the stronger relationship that we observed for photosynthesis.

However, our data indicate that communities with high growing season productivity likely also suffer higher yearly soil carbon losses, as we found no clear links with plant height and soil organic carbon stocks, either in the organic or the entire soil profile, that reflect the long-term annual carbon accumulation. This was an interesting finding because we hypothesized the relationship between plant size and soil organic carbon stocks to be positive as larger communities might produce larger carbon inputs to soils (DeMarco, Mack, and Bret-Harte 2014). The lack of this relationship suggests that in this ecosystem, other mechanisms have a stronger control over soil organic carbon stocks. Such mechanisms can be linked to carbon cycling processes during the shoulder and winter seasons or lateral transport of carbon, for example. One such mechanism is that during the winter, taller communities have warmer soils (Kropp et al. 2020), which accelerate soil respiration and increase soil carbon losses (Natali et al. 2019). Or, tall plant communities might inhabit environments where carbon leaching is high (Ma et al. 2019). In any case, our results show that tundra soil organic carbon stocks might be weakly linked to changes in growing season ecosystem CO_2_ fluxes, above-ground carbon stocks, and plant height, similar to the findings by Sørensen et al. (2018).

Evidence suggests that plant height is increasing around the Arctic in response to climate change. Sometimes this is in part due to “shrubification”, the increase in the biomass, cover, and abundance of woody species (Myers-Smith et al. 2011; Berner et al. 2020), but the height of non-woody species is increasing as well. Bjorkman et al. (2018) reported an average increase of about 1 cm in plant height over 1989–2015 years across the tundra. Our results suggest that such an increase would have resulted in 8–16% larger growing season photosynthesis and 6–11% larger ecosystem respiration (assuming a linear response to height, 90% CI). Our observational study thus indicates that the documented changes in plant height across the tundra might have already caused large shifts in the functioning of Arctic ecosystems, especially on growing season CO_2_ fluxes and above-ground carbon storage, but the effects on total ecosystem carbon storage require further investigation. New measurements in taller plant communities are likely important to address this as our study design did not cover the full plant height spectrum observed across the tundra; community-weighted plant height varied between 1 and 20 cm in this study whereas it varied between 0 and 40 cm in Bjorkman et al. (2018). In particular, our study design lacked tall Salix-dominated communities which are common in many southern parts of the Arctic, and are also one of the main drivers of tundra shrubification (Myers-Smith et al. 2011).

Earlier studies in the tundra have often characterized the effects of vegetation functional composition on carbon cycling with LAI (Street et al. 2007; Shaver et al. 2007; Dagg and Lafleur 2011; Marushchak et al. 2013). Measuring LAI does not require information on species assemblages, making it useful for landscape-scale or temporal predictions of carbon cycling (Shaver et al. 2007; Cahoon, Sullivan, and Post 2016). However, LAI is affected by several individual traits as, for example, increases in plant height and SLA can both increase LAI (Fletcher et al. 2012). Consequently, understanding the mechanisms driving the relationships between the environment and LAI, or LAI and carbon cycling, can be challenging. Our study adds to the results of these earlier studies by describing community functional composition with three widely measured above-ground plant traits that are related to fundamental trade-offs in ecological strategies, thus improving our mechanistic understanding about vegetation-carbon linkages in the tundra.

### Communities with fast traits are associated with higher soil organic carbon stocks

The effects of SLA and LDMC on carbon cycling were more variable and in general weaker compared to the relationships between plant height and carbon cycling. Our study supports the well-established strong positive relationship between faster economic traits (i.e. higher SLA and lower LDMC) and photosynthesis (Williams et al. 2006; Street et al. 2007; Shaver et al. 2007), but suggests that the connection to ecosystem respiration is not as strong. The relationship between ecosystem respiration and economic traits is more complex due to the various sources of ecosystem respiration (above-ground plant tissues, roots, symbionts, and heterotrophs) which all have different drivers (Segal and Sullivan 2014; Barba et al. 2018). We speculate that there are four potential mechanisms that explain higher ecosystem respiration associated with faster communities. First, higher levels of photosynthesis of fast communities also accelerates plant respiration (Lund et al. 2010). Second, faster-growing plants often produce more litter that is rich in nutrients and more easily broken down by soil microbes, resulting in higher soil respiration rates (Cornwell et al. 2008; G. T. Freschet, Aerts, and Cornelissen 2012). Third, faster plants produce more root exudates, which are, together with root respiration, suggested to be a main source of soil respiration in the tundra (Illeris, Michelsen, and Jonasson 2003). And fourth, plants with high leaf nutrient concentrations also have high nutrient concentrations in the roots, as suggested by the whole-plant economic spectrum hypothesis (G. T. Freschet et al. 2010), which might also accelerate soil respiration by facilitating root respiration and root litter decomposition (S. Jia et al. 2013). Thus, there are several processes that could be associated with the relationships that we have observed, all consistent with our observation that fast leaf economic traits accelerate both respiratory and photosynthetic fluxes.

Even though plant height did not have a clear effect on soil organic carbon stocks, SLA and LDMC had a relatively strong relationship with it. This might indicate that differences in organic matter quality (associated with leaf economics) instead of quantity (associated with larger plants) controls soil organic carbon stocks in this landscape (Hobbie 1996). Similar to Sørensen et al. (2018), largest soil organic carbon stocks in our study were located in graminoid and forb-dominated meadow communities with fast traits whereas smallest soil organic carbon stocks were found in communities with high LDMC and low SLA. Therefore, although it is generally hypothesized that fast communities often produce litter that decomposes rapidly, and that slow communities associated with evergreen species produce recalcitrant litter that decomposes slowly and accumulates into the soil (Hobbie et al. 2000), these hypotheses did not explain our results. Instead, our results suggest that mechanisms related to roots or microorganisms likely have a stronger control on soil organic carbon stocks. For example, communities with fast traits might have deeper-reaching roots and larger amounts of biomass below-ground (Ylänne et al. 2018; Iversen et al. 2015), and conditions deeper in the soil might make root litter relatively resistant to microbial decomposition compared to leaf litter and root litter in higher soil layers (De Deyn, Cornelissen, and Bardgett 2008; G. T. Freschet et al. 2013). Further, as speculated by Sørensen et al. (2018), microorganisms could retain more carbon in the soil due to the arbuscular mycorrhiza associated with faster communities whereas slower communities with ecto- and ericoid mycorrhiza could decompose organic matter faster (Väre, Vestberg, and Eurola 1992; Becklin, Pallo, and Galen 2012; Parker, Subke, and Wookey 2015). Moreover, communities with slow leaf economics often have evergreen leaves that produce relatively small carbon inputs to the soil compared to faster communities that lose their leaves each year (Hobbie 1996).

Nevertheless, a large part of variation in soil organic carbon stocks remained unexplained, even when soil moisture and air temperature conditions were considered. It is possible that these weaker links are partly due to practical reasons, as we measured soil organic carbon outside the vegetation collar (see Fig. S1). However, our study design contained so much variation in community-weighted SLA and LDMC that it is in this sense representative of almost the entire tundra biome (Thomas et al. 2020), thus our results should provide relatively robust results describing the direction and magnitude of the relationship between leaf economic traits and carbon cycle variables.

### Functional diversity in SLA and LDMC increases ecosystem functioning

All fluxes and soil organic carbon stocks correlated positively with the functional diversity in SLA and LDMC, but not with the diversity in plant height. Thus, within-community variability in leaf traits related to the leaf economics spectrum increased ecosystem functioning, supporting the niche complementarity hypothesis (Cadotte, Carscadden, and Mirotchnick 2011). Similar findings of a positive correlation with plant diversity and soil organic carbon stocks have also been made in earlier observational (Chen et al. 2018) and experimental (Lange et al. 2015; Fornara and Tilman 2008) studies. There are several mechanisms for how soil organic carbon stocks could increase with plant functional diversity. One such mechanism is that increased productivity could lead to higher litter inputs from leaves and roots (DeMarco et al. 2014; Chen et al. 2018). This is plausible, since leaf economic diversity also increases photosynthesis and the ratio of photosynthesis to ecosystem respiration. Niche complementarity pertaining to nutrient-use strategies could also lead to more efficient use of below-ground niche space and thus higher total investments in acquiring below-ground resources. Other mechanisms are related to soil microbes. High plant diversity might enhance the diversity and activity of soil microbial communities and decrease carbon losses from microbial decomposition (Chen et al. 2018).

Functional diversity in plant height was positively if somewhat uncertainly associated with above-ground carbon stocks, indicating that multi-layered vegetation stores more carbon compared to less structurally diverse plant communities. Similar effects of size-related trait diversity on above-ground biomass have been reported for temperate semi-natural grasslands (Schumacher and Roscher 2009). On the other hand, Conti and Díaz (2013) reported a negative correlation between structural diversity and above-ground carbon stocks in subtropical forests, but note that this is probably due to confounding factors. Our results provide more evidence that structurally layered plant communities might contain more carbon than vertically homogeneous vegetation patches, although the effects were uncertain enough that further research is needed to verify this relationship.

### Characterizing the underground community should be a priority

Our results showed that average plant height, SLA, and LDMC, and within-community variability in these traits were good predictors of above-ground carbon stocks and ecosystem fluxes whereas a large part of variation in soil organic carbon stocks remained unexplained. This is an issue as most of the carbon in tundra ecosystems is located in the soil (Hugelius et al. 2014; Mishra et al. 2021). Field and laboratory evidence suggests that this carbon pool is vulnerable to warming and permafrost thaw (Voigt et al. 2017; Schädel et al. 2016), but it is not well known how it responds to the wide-spread shifts in tundra vegetation (Bjorkman et al. 2018; Berner et al. 2020; Parker et al. 2021). This is because soil organic carbon stocks are a result of various carbon inputs from above- and below-ground plant tissues, outputs from ecosystem and soil respiration, and lateral transport, which all respond differently to trait variation. Therefore, interpreting the effects of above-ground traits on soil organic carbon stocks is challenging as the links might be weak and indirect (e.g., via litter decomposition and accumulation). Further, soil organic carbon stocks are strongly driven by abiotic environmental conditions as well, some of which might have been missing from our model. For example, soil organic carbon stocks can be influenced by geomorphological processes such as cryoturbation, or aeolian and fluvial transportation of litter (Klaminder, Yoo, and Giesler 2009), or soil development and carbon accumulation in the past (Palmtag et al. 2015).

Most importantly, our results highlight that we need more measurements and understanding on belowground community composition, diversity and functioning of plant roots, litter, and microorganisms (Orwin et al. 2010). With recent advancements such as a new a root ecology handbook (G. Freschet et al. 2020) and the identification of major axes of variation in below-ground traits (Bergmann et al. 2020), some of these measurements can soon be done in a comparable way across the globe. Such measurements will help us understand the mechanisms influencing soil organic carbon stocks and vegetation-carbon feedbacks in a changing climate.

## Conclusions

We found that carbon cycling in the tundra was well explained by plant height, SLA, and LDMC, and their within-community variabilities. Average plant height was the strongest predictor of most carbon cycling variables. Diversity in SLA and LDMC mattered strongly for soil organic carbon stocks, highlighting a potentially important mechanism controlling the vast carbon pools that should be better recognized. Plant height increased growing season CO_2_ fluxes and above-ground carbon stocks but had no clear effect on soil organic carbon stocks, whereas fast leaf economics were associated with higher CO_2_ fluxes and soil organic carbon stocks. By utilizing globally applicable plant functional traits, our findings facilitate forming an integrated view on the causes and ecosystem consequences of climate change-related vegetation transitions in the tundra.

## Acknowledgements

The authors thank Liangzhi Chen, Elisa Hanhirova, Arttu Kivimäki, Panu Lammi, Meri Lindholm, Mitro Müller, Tiia Määttä, Aino-Maija Määttänen, Annina Niskanen, Nina Nordblad, Helena Rautakoski, Henri Riihimäki, Tuuli Rissanen, Sakari Sarjakoski, Outi Seppälä, Akseli Toikka, Vilna Tyystjärvi, and the Arctic Microbial Ecology group at the University of Helsinki for their field and laboratory work, and the staff of the Kilpisjärvi Biological Station for their help. We also thank Markus Hartman at Natural Resources Institute Finland for borrowing measurement equipment. KH was funded by the Doctoral Programme in Wildlife Biology Research at the University of Helsinki. A-MV was funded by Societas pro Fauna et Flora Fennica, Otto Malm foundation, Nordenskiöld-samfundet, Alfred Kordelin foundation, Finnish Cultural Foundation, Academy of Finland (project no. 286950), and Väisälä fund. JK was funded by the Doctoral Programme in Geosciences at the University of Helsinki, Maaja vesitekniikan tuki ry. and Tiina and Antti Herlin Foundation. The authors acknowledge funding from the Academy of Finland (project 286950).

