## Appendix S1 for "Relationships between aboveground plant traits and carbon cycling in tundra plant communities"

The following Supporting Information is available for this article:

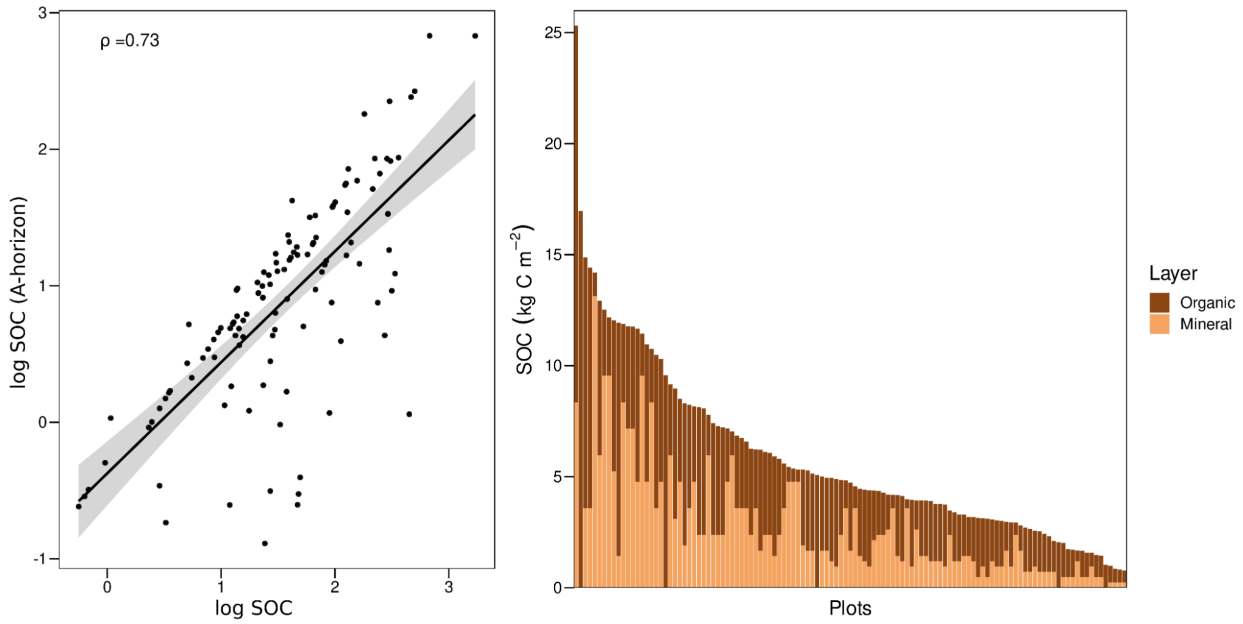


Fig. S1. Relationship between Total soil organic carbon (SOC) stocks and organic layer (A-horizon) SOC stocks.

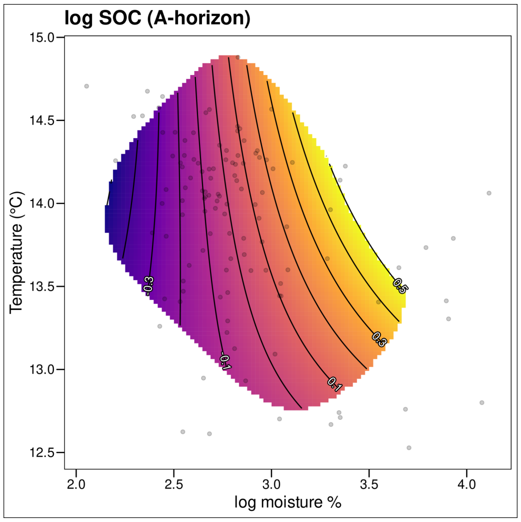


Fig. S2. Marginal effects of 3-year average July air temperature and growing-season soil moisture on organic layer SOC stocks. Point clouds show the observed data, and contour lines represent predicted relative responses on the log-scale. Responses are shown only where the standard error of the smoother is less than ¼ of the observed standard deviation.

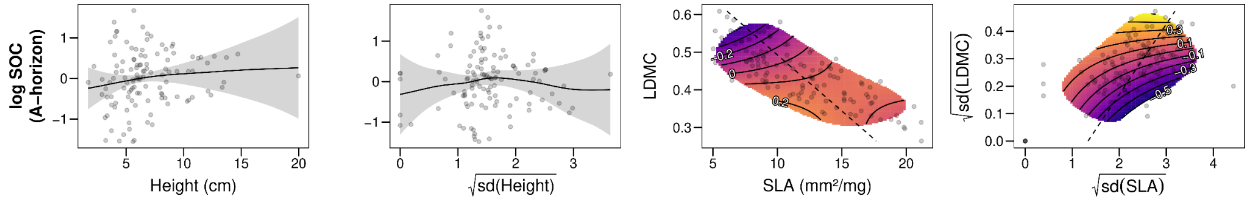


Fig. S3. Marginal responses of organic layer soil organic carbon stocks to plant height, SLA, LDMC and their within-community variability. In the two leftmost panels, the shading represents two standard errors and the points are partial residuals. In the two rightmost panels, point clouds show the observed data and contour lines represent predicted relative responses on the log-scale. The 2D response surfaces are limited to where the standard error of the smoother is less than ¼ of the observed standard deviation. A dashed line shows the location of a model II regression line depicting the direction of covariation between SLA and LDMC, and their variabilities.

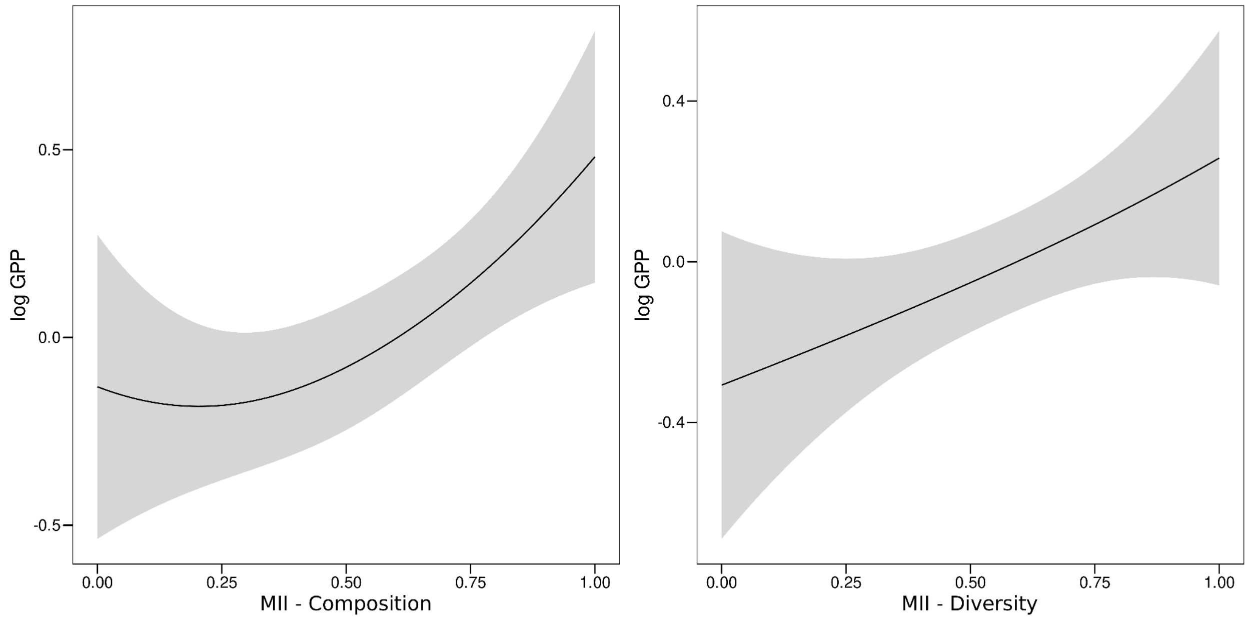


Fig. S4. Response of gross primary productivity (GPP) to the model II regression line between LDMC and SLA (MII - Composition), and their variabilities (MII - Diversity). Higher values of the compositional axis mean faster leaf traits i.e. higher SLA and lower LDMC. The lines correspond to the dashed lines in Figure 3. The shaded area is a credible interval of two standard errors.

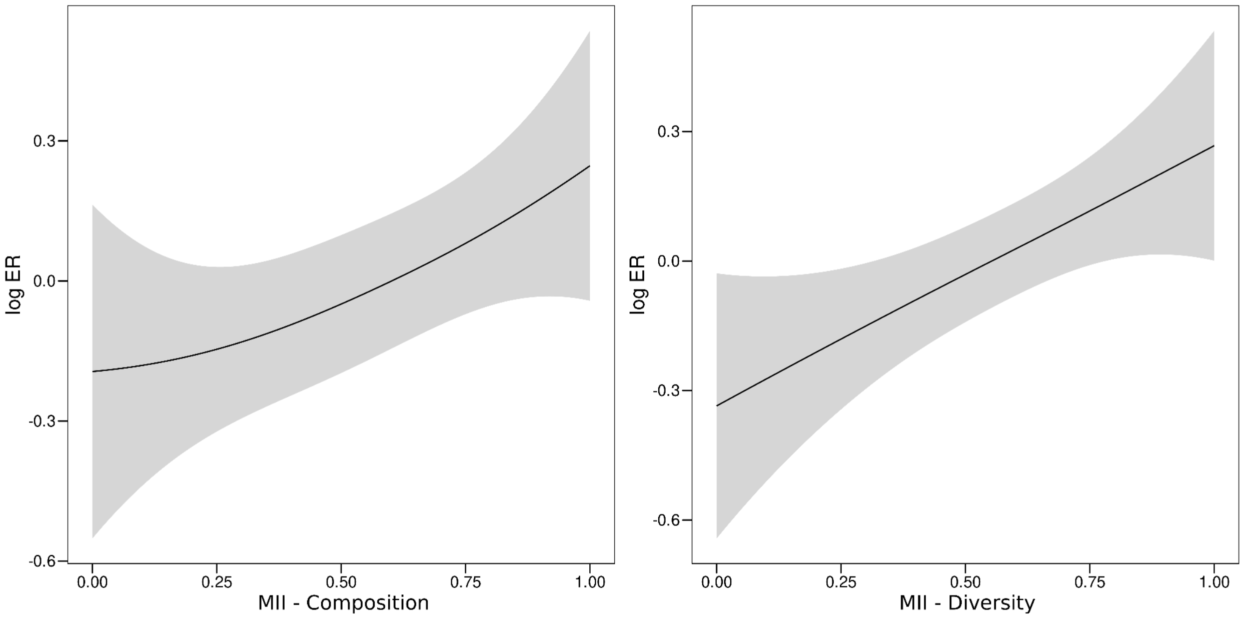


Fig S5. Response of ecosystem respiration (ER) to the model II regression line between LDMC and SLA (MII - Composition), and their variabilities (MII - Diversity). Higher values of the compositional axis mean faster traits i.e. higher SLA and lower LDMC. The lines correspond to the dashed lines in Figure 3. The shaded area is a credible interval of two standard errors.

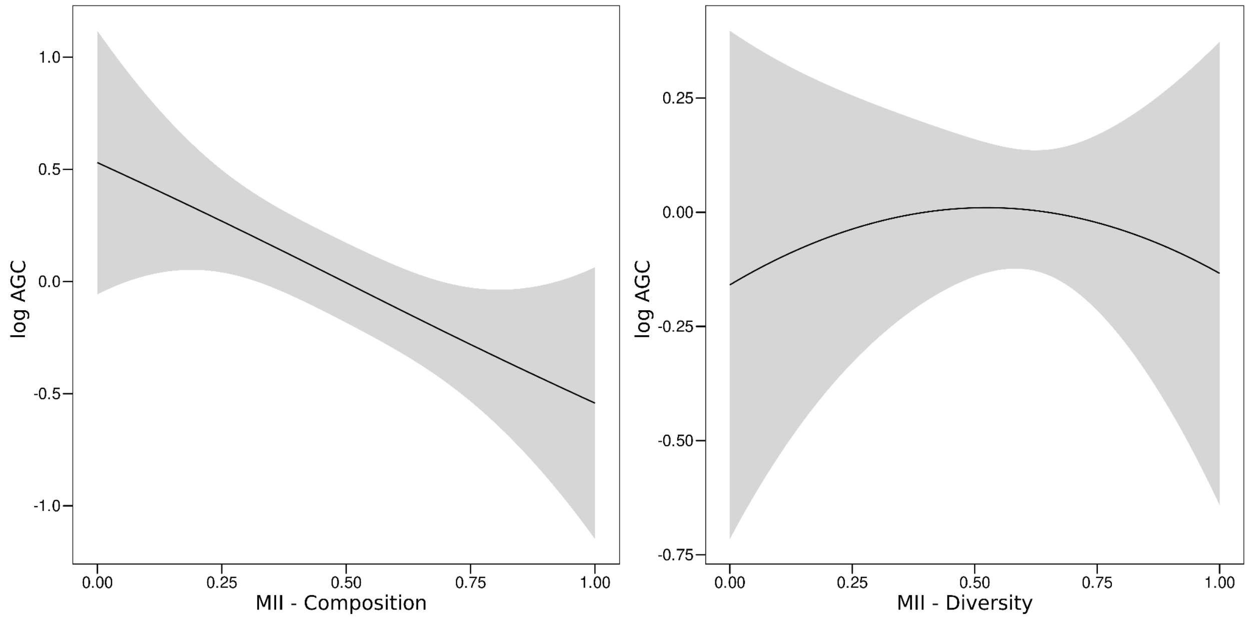


Fig. S6. Response of above-ground carbon stocks (AGC) to the model II regression line between LDMC and SLA (MII - Composition), and their variabilities (MII - Diversity). Higher values of the compositional axis mean faster traits i.e. higher SLA and lower LDMC. The lines correspond to the dashed lines in Figure 3. The shaded area is a credible interval of two standard errors.

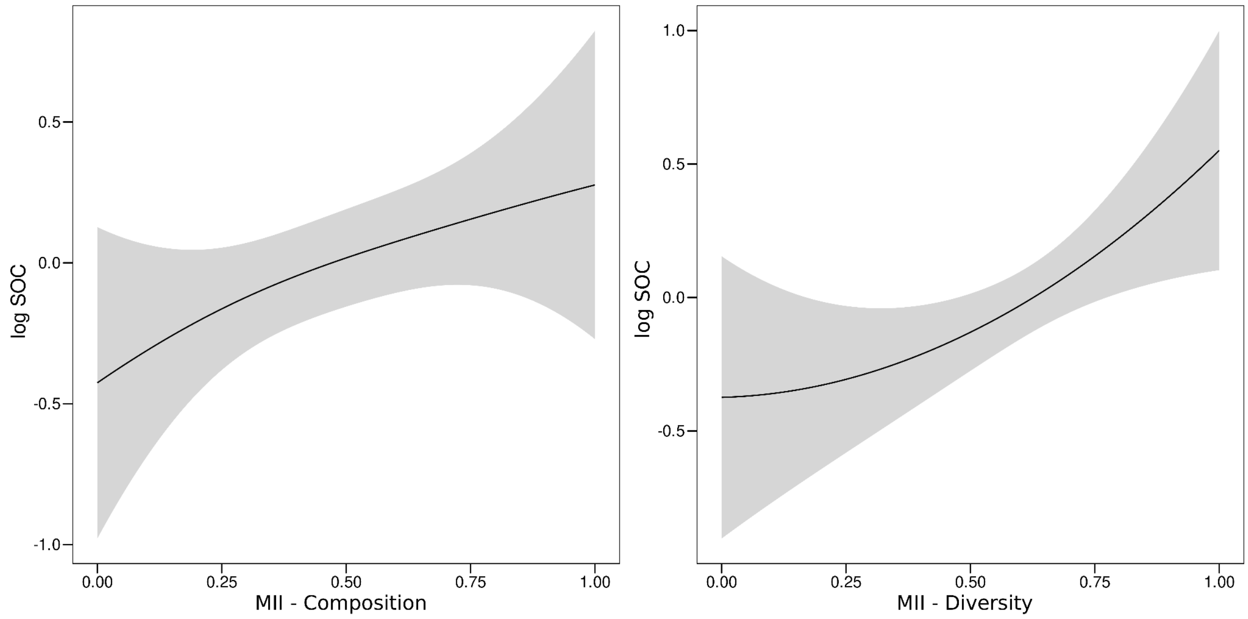


Fig. S7. Response of soil organic carbon stocks (SOC) to the model II regression line between LDMC and SLA (MII - Composition), and their variabilities (MII - Diversity). Higher values of the compositional axis mean faster traits i.e. higher SLA and lower LDMC. The lines correspond to the dashed lines in Figure 3. The shaded area is a credible interval of two standard errors.

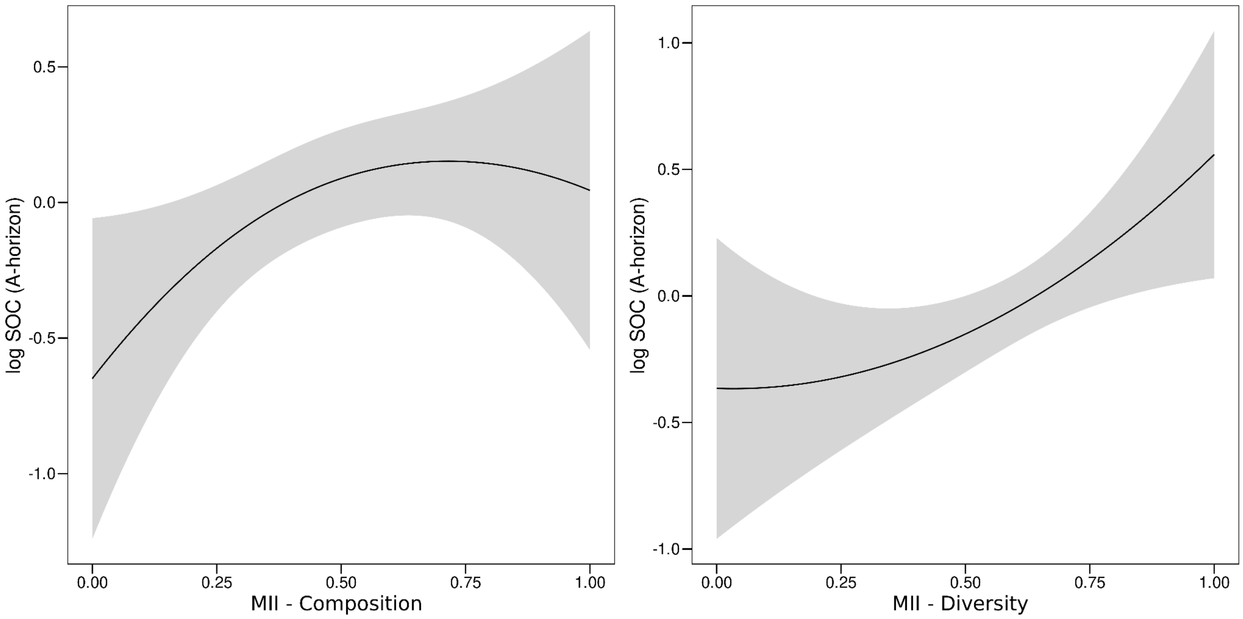


Fig. S8. Response of organic layer soil organic carbon stocks (SOC, A-horizon) to the model II regression line between LDMC and SLA (MII - Composition), and their variabilities (MII - Diversity). Higher values of the compositional axis mean faster traits i.e. higher SLA and lower LDMC. The lines correspond to the dashed lines in Figure 3. The shaded area is a credible interval of two standard errors.

**Table S1.** Details on the measurement equipment.

| **Variable** | **Description of the measurement device** |
|---|---|
| **Abiotic variables** | |
| Soil moisture | A handheld time-domain reflectometry sensor of the accuracy ±3.0 VWC% with 0.1 VWC% resolution (FieldScout TDR 300; Spectrum Technologies Inc., Plainfield, IL, USA) |
| Air temperature | Thermochron iButton DS1921G and DS1922L with a temperature range between -40 and 85 °C, resolution of 0.5°C, and accuracy of 0.5°C |
| **Carbon cycle variables** | |
| Carbon dioxide probe | GMP343 A1A1N0N0N0A model with $CO_2$ range 0–1000 ppm, ± 3 ppm + 1% (Vaisala, Vantaa, Finland). Concentrations of $CO_2$ were already corrected for atmospheric pressure and relative humidity during the measurements. |

| | |
|---|---|
| Air humidity and air temperature probe HMP75 | HMP75 with a temperature range between -20 and 60°C, relative humidity range 0–100 %; temperature accuracy at 20°C ±0.2°C, and relative humidity accuracy between -20 and + 40°C ± (1.0 + 0.008 * relative humidity reading)) (Vaisala, Vantaa, Finland). |
| Measurement indicator MI70 | Data reader for GMP343 and HMP75 (Vaisala, Vantaa, Finland). |
| Quantum sensor with a hand-held meter | MQ-200 quantum sensor with a hand-held meter (Apogee Instruments, Inc, USA). MQ-200 measures PAR at a spectral range from 410 to 655 nm. |
| C% | Total carbon content (C%) analyses were done using either Vario Elementar Micro cube or Vario Elemantar Max -analyzer (Elementar Analysensysteme GmbH, Germany). The laboratory analyses were carried out in the Laboratory of Geosciences and Geography and Laboratory of Forest Sciences (University of Helsinki). |
